# *In Vitro* Maturation of Bone Marrow-Derived Dendritic Cells via STING Activation for T Cell Priming

**DOI:** 10.1101/2025.06.11.659197

**Authors:** Busra Buyuk, Kaiming Ye

## Abstract

Dendritic cells (DCs) are among the most potent antigen-presenting cells, playing a pivotal role in initiating adaptive immune responses. The STING (Stimulator of Interferon Genes) pathway, a key cytosolic DNA-sensing mechanism, has emerged as a powerful modulator of type I interferon production and CD8⁺ T cell activation. In this study, we aimed to generate and functionally mature bone marrow-derived DCs *in vitro* by optimizing cytokine conditions and incorporating STING pathway stimulation. Immature DCs were differentiated from murine bone marrow using various concentrations of GM-CSF and IL-4, and characterized by the expression of surface markers CD11c, CD80, and MHC-II. Maturation was induced using a STING agonist, followed by co-culture with naïve CD8⁺ T cells isolated from mouse spleens via Magnetic Activated Cell Sorting (MACS). STING-activated DCs exhibited enhanced surface marker expression and significantly promoted CD8⁺ T cell proliferation. Our findings demonstrate that combining optimized cytokine-driven differentiation with STING activation significantly improves DC immunogenicity, offering a promising platform for the development of DC-based cancer immunotherapies.

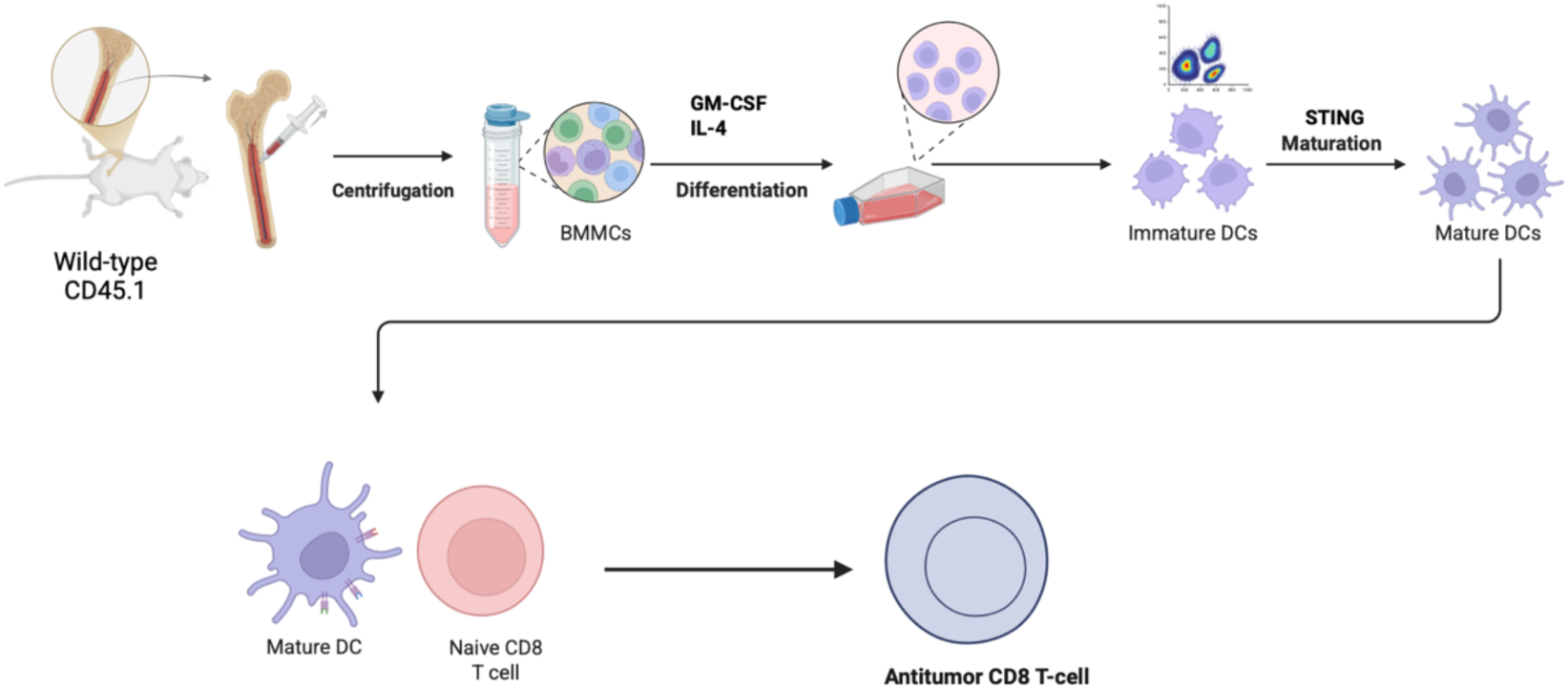

Graphical abstract created using BioRender.com

## 1. Introduction

Dendritic cells (DCs) play a critical role in the innate immune system^1, 2^. They serve as the most potent antigen-presenting cells that bridge the innate and adaptive immune systems^3, 4^. Their ability to capture, process, and present antigens to T cells plays a central role in initiating immune responses that include CD8+ T cells, which are essential for fighting viral infections and cancer^4^. Differentiation of DCs from hematopoietic stem cells in the bone marrow can be induced *in vitro* using cytokines such as Granulocyte-Macrophage Colony-Stimulating Factor (GM-CSF) and Interleukin-4 (IL-4)^5, 6^. These cytokines play an important role in immature DC maturation by enhancing their capacity to present antigen and activate T cells^7^. The DC differentiation and maturation process is regulated by several factors, including the expression of surface markers such as CD11, CD80, and MHC II ^8, 9^. These markers, along with other markers, can help measure the antigen-presenting cell capacity of DCs. GM-CSF and IL-4 are widely used to induce DC differentiation, but the optimal concentrations to achieve the highest level of antigen-presenting capacity of DCs remain unclear ^8^. A better understanding of how these cytokines influence DC maturation could enhance the use of DCs in immunotherapy applications, such as cancer treatment, by improving their ability to activate immune cells effectively ^10^.

The STING pathway is important in regulating immune responses ^11^. STING is a cytosolic receptor that senses pathogenic DNA in the cell and initiates a signaling cascade leading to the production of type I interferons ^11, 12^. These interferons activate immune cells, including DCs, and promote the development of an inflammatory environment that enhances immune responses^13^. DCs enhance the antigen-presenting function of STING activation and promote the activation of CD8+ T cells, which are important for immune defense against pathogens and tumors^14^. Although the effects of STING activation on DC function are well known, the optimal conditions for combining STING activation with DC differentiation have not been fully explored^15^. This study aims to investigate the role of different GM-CSF and IL-4 concentrations in DC differentiation, focusing on the expression of key surface markers (CD11c, CD80, MHC II). It also examines how STING pathway activation influences DC maturation and functional activation potential^16^. In this study, we used bone marrow-derived DCs induced with GM-CSF and IL-4 and then exposed them to different concentrations of STING agonist. We evaluated the expression of CD11c, CD80 and MHC II in these DCs and evaluated their ability to induce CD8 T cytotoxic activation and proliferation by co-culturing mature DCs with naive CD8+ T cells. Our goal was to optimize conditions that best support *in vitro* priming of naive T cells, by identifying cytokine and STING agonist combinations that promote robust DC function.

## 2. Materials and Methods

### 2.1 Dendritic Cell (DC) Isolation from Mouse Bone Marrow

#### 2.1.1 Bone Marrow Extraction

All animal procedures were performed in accordance with an IACUC-approved protocol at Binghamton University, following guidelines from the American Veterinary Medical Association. Eight C57BL/6 mice (Jackson Laboratory) were euthanized via CO₂ inhalation. The fur was removed and the legs were disinfected with 70% ethanol. Skin and muscle tissues were carefully dissected using sterile surgical scissors and forceps to expose the femurs and tibias, as well as front leg bones. These bones were placed in 50 mL conical tubes containing cold Isolation Buffer (1X DPBS, 2% FBS, 2 mM EDTA) and transferred to a BSL-2 laminar flow cabinet for further processing. Connective tissues were removed, and the cleaned bones were placed in sterile, non-treated petri dishes containing isolation buffer. Bone marrow was flushed using a 1 mL syringe with a 27 G needle into 100 mm non-treated petri dishes with DC culture medium (RPMI-1640, 10% heat-inactivated FBS, 2 mM L-glutamine, 50 μM β-mercaptoethanol, 1X penicillin/streptomycin), excluding GM-CSF and IL-4. Each bone was flushed multiple times until it appeared white, ensuring complete marrow removal. The resulting suspension was pipetted to disperse cell clumps and filtered through a 70 μm cell strainer into a 50 mL conical tube. The remaining marrow was rinsed with medium and filtered again. Cells were centrifuged at 1,500 rpm for 8 minutes. The pellet was resuspended in 3 mL RBC Lysis Buffer (150 mM NH₄Cl, 10 mM KHCO₃, 0.1 mM EDTA, pH 7.2–7.4) and incubated on ice for 5 minutes. Lysis was stopped by adding 20 mL DPBS followed by centrifugation at 1,500 rpm for 8 minutes. The pellet was resuspended in 2 mL of DC culture medium and counted using trypan blue exclusion and a hemocytometer under a 10x phase-contrast inverted microscope. Cells were seeded equally into four 100 mm tissue culture-treated dishes containing 10 mL of DC culture medium supplemented with 20 ng/mL GM-CSF and 20 ng/mL IL-4.

#### 2.1.2. DC Culture

Cells were incubated at 37°C in a humidified incubator with 5% CO₂. On Day 3, cultures were examined using an inverted phase-contrast microscope to assess morphology and cell density. On Day 6, non-adherent and loosely adherent cells, enriched for immature dendritic cells, were collected by gentle pipetting, centrifuged (1,500 rpm, 8 min), and resuspended in fresh dendritic cell (DC) culture medium. During medium change steps, only the floating (non-adherent) cells were collected; the adherent cells that remained attached to the plate likely macrophage-rich were left behind. In selected experiments, the concentrations of granulocyte-macrophage colony-stimulating factor (GM-CSF) and interleukin-4 (IL-4) were varied to optimize dendritic cell yield and phenotype. DC Culture Medium Composition: RPMI-1640 supplemented with 10% heat-inactivated fetal bovine serum (HI-FBS), 2 mM L-glutamine, 50 μM β-mercaptoethanol, 1× penicillin-streptomycin, 20 ng/mL GM-CSF, and 20 ng/mL IL-4.

#### 2.1.3. Dendritic Cell Differentiation and Cytokine Optimization

Bone marrow cells were plated into 100 mm non-treated culture dishes at a density of 1 × 10⁶ cells/mL in 10 mL of complete RPMI 1640 medium and cultured at 37°C in a humidified 5% CO₂ incubator. To determine optimal cytokine conditions for dendritic cell (DC) generation, cells were cultured for 10 days under four different media supplementation. To optimize cytokine conditions for dendritic cell (DC) differentiation, four different media supplementation protocols were tested over a 10-day culture period. In the first condition, 20 ng/mL of GM-CSF was added on days 0, 3, 6, and 8. In the second condition, cells were treated with 20 ng/mL GM-CSF on days 0 and 3, followed by 10 ng/mL GM-CSF and 10 ng/mL IL-4 on days 6 and 8. In the third condition, 20 ng/mL GM-CSF was added on days 0 and 3, then reduced to 10 ng/mL GM-CSF on days 6 and 8. In the fourth condition, GM-CSF was added in decreasing concentrations: 20 ng/mL on day 0, 10 ng/mL on day 3, 5 ng/mL on day 6, and 2.5 ng/mL on day 8. At each time point, the culture medium was completely removed and replaced with 10 mL of fresh medium containing the respective cytokine concentrations. During each medium change, only the floating cells in the supernatant were collected and centrifuged; the adherent cells remaining in the plate were not included, as they were presumed to be macrophage-enriched. Cells were harvested after on day 10 for downstream analysis.

### 2.2. Immunostaining

For flow cytometric analysis, 1×10⁶ cells in 100 µL medium were transferred into wells of a 96-well U-bottom non-tissue culture-treated plate (Corning). Cells were centrifuged at 1,500 rpm for 8 min and resuspended in 100 µL of Fc block (CD16/CD32, BioLegend) diluted 1:100 in FACS buffer (1X DPBS, 25 mM HEPES, 2 mM EDTA, 2% FBS). The plate was incubated on ice for 15 min. Antibody cocktails for surface markers (CD11c/CD80 and CD11c/MHC II) were prepared in FACS buffer. CD11c and CD80 antibodies were diluted 1:100, while MHC II was diluted 1:300. 100 µL of each antibody cocktail was added to corresponding wells, followed by a 30-minute incubation on ice protected from light. Cells were washed with 150 µL FACS buffer, centrifuged, and fixed with 100 µL 4% paraformaldehyde (PFA) at room temperature for 10 minutes in the dark. After fixation, cells were washed, resuspended in 100 µL FACS buffer, and transferred to FACS tubes containing 200 µL FACS buffer for analysis.

#### 2.2.1. STING Stimulation of Bone Marrow-Derived Dendritic Cells

To investigate the effect of STING activation on the maturation of bone marrow-derived dendritic cells (BMDCs), cells cultured under optimized conditions (GM-CSF + IL-4) were stimulated on day 9 with a STING agonist. The agonist used was 2′3′-c-di-AM(PS)₂ (Rp,Rp) VacciGrade™ (InvivoGen), a synthetic cyclic dinucleotide known to specifically activate the STING pathway. BMDCs were maintained in media containing 10 ng/mL GM-CSF and 10 ng/mL IL-4 and transferred to 6-well plates on day 9. Cells were treated with the STING agonist at three different final concentrations: 2.5 µg/mL, 5 µg/mL, and 20 ng/mL (low dosoge), in fresh complete RPMI 1640 medium supplemented with cytokines. Control wells received no STING agonist. All conditions were incubated for 24 hours at 37°C with 5% CO₂.

After 24 hours of stimulation, on day 10, cells were harvested and analyzed by flow cytometry for the expression of maturation markers CD80 and MHC II. The aim was to assess the effect of STING pathway activation on further promoting DC maturation under cytokine-supported conditions.

### 2.3. Dendritic Cell Characterization

#### 2.3.1 Flow Cytometry

Flow cytometric analysis was conducted at the Analytical and Diagnostics Laboratory (ADL) at Binghamton University. The antibodies and fluorophores used are detailed in **Table 1**.

**Table 1.** Antibodies for DC and T Cell Characterization.

| **Antibody** | **Target** | **Vendor** | **Catalog No.** | **Dilution** | **Fluorophore** | **Laser (Ex/Em)** |
| --- | --- | --- | --- | --- | --- | --- |
| CD11c | DCs | BioLegend | 117309 | 1:100 | APC | 637 nm / 655–685 nm |
| CD80 | DCs | BioLegend | 104715 | 1:100 | Alexa Fluor 647 | 488 nm / 515–545 nm |
| MHC II | DCs | BioLegend | 107607 | 1:300 | PE | 488 nm / 483–493 nm |
| CD8 | T cells | BioLegend | 100711 | 1:100 | APC | 637 nm / 655–685 nm |
| CD69 | T cells | BioLegend | 104507 | 1:100 | PE | 488 nm / 515–545 nm |

### 2.4 Spleen Harvest and Digestion

#### 2.4.1 Spleen Harvest

Two C57BL/6 mice were euthanized via CO₂ inhalation. After shaving and sterilization with 70% ethanol, the spleens were surgically removed using sterile scissors and forceps and placed in Basic Medium (RPMI-1640, 2% FBS, 1X P/S).

#### 2.4.2 Spleen Digestion

Spleens were minced in cold basic medium using sterile blades and transferred into 15 mL tubes containing 2 mL of digest buffer (RPMI-1640, 2% FBS, 1X P/S, 1 mg/mL Collagenase IV, 100 µg/mL DNase I). Tubes were incubated at 37°C for 30 min, vortexed every 5 min, and filtered through a 70 µm strainer. Cells were centrifuged at 1,500 rpm for 5 min, treated with RBC Lysis Buffer on ice for 5 min, then washed with 20 mL DPBS and centrifuged again.

#### 2.4.3 Cell Counting and Staining

Cells were counted using trypan blue exclusion. Concentration was calculated using the formula: Cells/mL = 10⁴ × 2 (dilution factor) × average cell count per square / 4

### 2.5 Magnetic Cell Sorting (MACS) for CD8⁺ T Cells

MACS was used to isolate CD8⁺ naïve T cells from digested spleens using BioLegend’s MojoSort™ system. Briefly, 10⁷ cells were incubated with 10 µL biotin-conjugated CD8 antibody cocktails for 15 min on ice, followed by 10 µL streptavidin-coated nanobeads for another 15 min. After incubation, 2.5 mL FACS buffer was added and tubes were placed in a compatible magnetic separator for 5 min. Unbound cells were collected and cultured in T cell medium for 3 days.

#### 2.5.1. T cell co-culture with dendritic cells

Naïve CD8⁺ T cells were co-cultured with dendritic cells (DCs) previously cultured in the presence of 10 ng/mL GM-CSF and 10 ng/mL IL-4, and stimulated with 5 µg/mL STING agonist. Co-culture was performed at a 1:5 DC-to-T cell ratio for 3 days in RPMI 1640 medium supplemented with 10% FBS and 1% penicillin-streptomycin.

### 2.6. CFSE Staining for T Cell Proliferation

CFSE labeling was performed to monitor T cell division. A total of 1×10⁶ T cells/mL were resuspended in 1 mL DPBS containing 0.5 µL CFSE (BioLegend, Cat. No. 423801). Cells were incubated at room temperature in the dark for 20 min. The reaction was quenched by adding 4 mL of T cell culture medium. Cells were centrifuged at 350 rpm for 5 min and resuspended for further assays or flow cytometry.

## 3. Results and Discussion

### 3.1. Optimization of a Culture Medium for Generating Bone Marrow-Derived Dendritic Cells

Dendritic cells (DCs) require growth factors for differentiation from bone marrow. In this study, we tested the effects of GM-CSF and IL-4 on DC production under four different culture conditions. Initially, GM-CSF was added fresh every three days, starting at a concentration of 20 ng/mL and gradually reducing to 2.5 ng/mL. In the second condition, a constant 10 ng/mL GM-CSF was used; in the third, 10 ng/mL GM-CSF was combined with 10 ng/mL IL-4; and the fourth condition consisted of 20 ng/mL GM-CSF only. On day 10, cells were analyzed by flow cytometry using antibodies against CD11c, CD80, and MHC II. The results showed that the third condition, with the combination of GM-CSF and IL-4, produced the highest proportion of CD11c-positive cells, with 84.7% of cells expressing CD11c Fig.1A. As shown in Fig.1B, DCs cultured with 10 ng/mL GM-CSF exhibited 27.4% MHC II-positive cells, while those cultured with 20 ng/mL GM-CSF showed 26.5%. In contrast, the combination of 10 ng/mL GM-CSF and 10 ng/mL IL-4 resulted in 34.1% MHC II-positive cells. These results indicate that IL-4 promotes MHC II expression and supports DC maturation. Next, the maturity of these cells was assessed by double staining for CD11c and CD80. As seen in Fig. 1C. CD80 expression decreased at lower GM-CSF concentrations and increased at higher concentrations. This suggests that GM-CSF enhances DC maturation.

**Figure 1.**
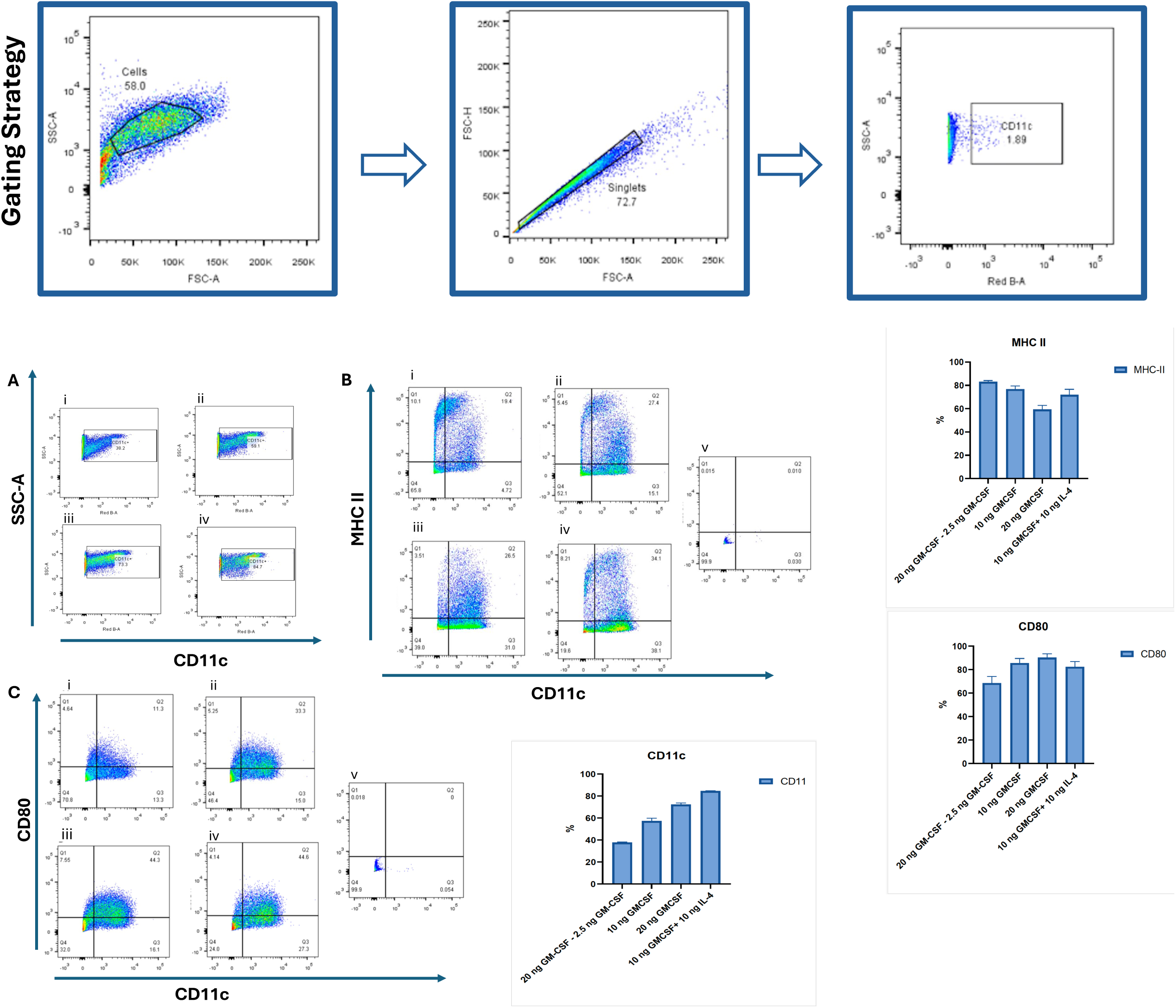
Phenotypic characterization of bone marrow-derived dendritic cells (BMDCs) cultured under different cytokine conditions. Bone marrow cells were cultured for 10 days in four different cytokine conditions to evaluate dendritic cell (DC) differentiation and maturation: (i) Condition 1 – Stepwise reduction of GM-CSF, starting at 20 ng/mL on day 0, then decreasing to 10 ng/mL (day 3), 5 ng/mL (day 6), and 2.5 ng/mL (day 8); (ii) Condition 2 –20 ng/mL GM-CSF (day 0, 3); 10 ng/ml GM-CSF (day 6, 8); (iii) Condition 3 –20 ng/mL GM-CSF; (day 0, 3, 6, 8); (iv) Condition 4 – 20 ng/mL GM-CSF (day 0, 3), followed by 10 ng/mL GM-CSF + 10 ng/mL IL-4 (day 6,8). Flow cytometry was performed on day 10 to assess the expression of dendritic cell surface markers. (A) CD11c expression as a general DC marker; (B) MHC class II expression, indicating antigen-presenting capability; (C) CD80 expression, representing DC maturation status. Unstained bone marrow cells were used as negative controls for gating. The data are presented as the mean ± SD of two technical replicates.

Overall, the data suggest that while high-dose GM-CSF increases the number of CD11c+ cells, it suppresses MHC II expression. In contrast, IL-4 helps to balance this suppression, promoting maturation. Therefore, the combination of GM-CSF and IL-4 provides the optimal conditions for generating both numerically abundant and immunologically mature DCs.

### 3.2. STING Agonist-Induced Maturation of Dendritic Cells

To assess whether STING pathway activation promotes dendritic cell (DC) maturation, bone marrow-derived DCs (BMDCs) were cultured for 9 days in the presence of 10 ng/mL GM-CSF and 10 ng/mL IL-4. On day 9, cells were stimulated with the STING agonist 2′3′-c-di-AM(PS)₂ (Rp,Rp) at varying concentrations (20 ng/mL, 2.5 µg/mL, and 5 µg/mL) and incubated for an additional 24 hours. 20 ng/mL dose was included as a low-range exposure, intended to evaluate whether minimal stimulation could initiate early activation markers. The 2.5 µg/mL and 5 µg/mL concentrations represent intermediate and high doses, respectively, based on prior literature for in vitro activation of mouse BMDCs. Untreated BMDCs served as negative controls. Flow cytometry was performed to evaluate the surface expression of maturation markers CD80 and MHC II. As shown in Fig.2A, CD80 expression was detected on 83.7% of untreated non-starved DCs. Upon stimulation with 20 ng/mL STING agonist, CD80 expression slightly increased to 88.8%. At higher doses of the STING agonist (2.5 µg/mL and 5 µg/mL), a noticeable increase in the median fluorescence intensity (MFI) of CD80 was observed Fig.2B, suggesting a dose-dependent upregulation of this maturation marker. However, the difference in CD80 MFI between the 2.5 µg/mL and 5 µg/mL doses was minimal, indicating a potential saturation effect. The modest difference in CD80 intensity at 20 ng/mL STING agonist compared to untreated DCs may reflect limited activation at low concentrations. We speculate that the continuous presence of GM-CSF and IL-4 throughout the culture period may influence the degree of CD80 upregulation upon STING stimulation. MHC II expression was also examined following STING agonist treatment.

**Figure 2.**
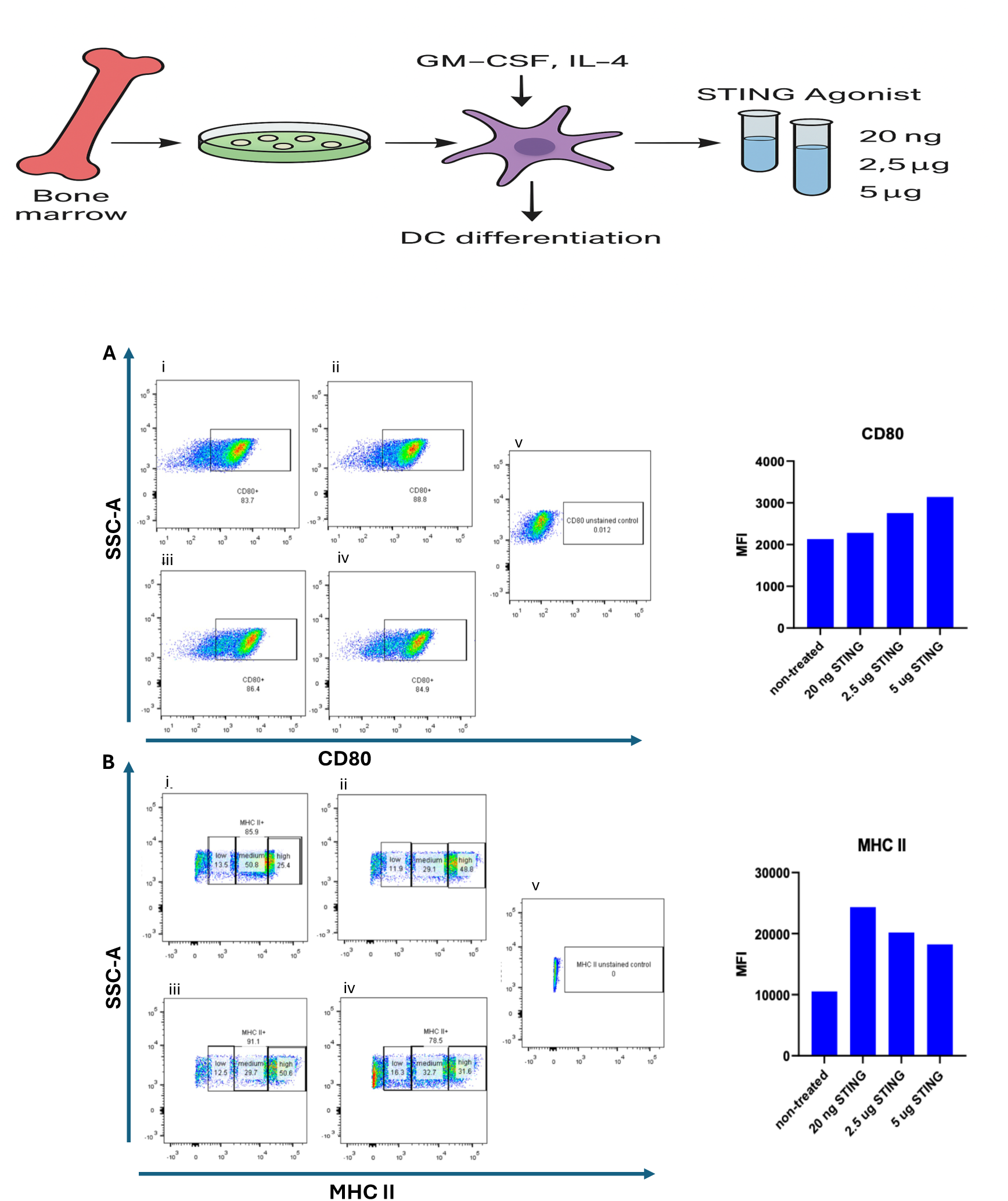
CD80 and MHC II expression in dendritic cells (DCs) cultured with GM-CSF and IL-4 and stimulated with a STING agonist. Bone marrow-derived dendritic cells (BMDCs) were cultured for 9 days in the presence of 10 ng/mL GM-CSF and 10 ng/mL IL-4. On day 9, cells were stimulated for 24 hours with increasing concentrations of the STING agonist 2′3′-c-di-AM(PS)₂ (Rp,Rp) at the following doses: 20 ng/mL, 2.5 µg/mL, and 5 µg/mL. Untreated cells served as negative controls, and unstained cells were included to establish flow cytometry gating. Flow cytometry was used to evaluate the surface expression of the DC maturation markers CD80 and MHC class II (MHC II). (A) Representative histograms showing CD80 expression in untreated and STING-stimulated DCs. (B) Representative histograms showing MHC II expression in untreated and STING-stimulated DCs. Median fluorescence intensity (MFI) of CD80 and MHC II expression across all treatment conditions. Data represent one independent sample per treatment group.

As depicted in Figure 2B, untreated DCs exhibited the lowest MFI of MHC II expression. Interestingly, stimulation with the lowest dose (20 ng/mL) of STING agonist resulted in the highest MFI for MHC II, while treatment with 2.5 µg/mL yielded a slightly lower, but still elevated, MFI. The highest dose (5 µg/mL) did not further enhance MHC II intensity and showed a marginal decrease compared to 2.5 µg/mL. These findings indicate that while STING activation can upregulate MHC II expression in DCs, a higher agonist concentration does not necessarily correlate with greater activation, possibly due to receptor saturation or negative feedback mechanisms.

### 3.3. CD8+ Naive T Cell Activation and Proliferation

Fig. 3 illustrates the gating strategy used to detect the activation and proliferation of CD8+ naïve T cells. After isolating these cells, they were stained with CD8 and CD69 antibodies to distinguish naïve CD8+ T cells and assess their activation status, respectively. Additionally, to monitor proliferation, the cells were labeled with CFSE dye on the first day of co-culturing and kept stained for three days. On the third day, activation was evaluated based on CD69 expression. In the absence of any stimulation, only 0.66% of the CD8+ naïve T cells showed activation. However, when these cells were co-cultured with unstimulated dendritic cells (DCs), the activation rate increased to 3.16%. Notably, a higher activation level of 9.03% was observed when CD8+ naïve T cells were co-cultured with dendritic cells that had been stimulated with 5 µg/mL of a STING agonist, indicating that this stimulation effectively triggers T cell activation. In terms of proliferation, a low level of cell division was detected in T cells co-cultured with unstimulated DCs. In contrast, a significant proliferation rate of 28.8 (15.7+ 13.1) % Fig.3B. was observed when CD8+ T cells were co-cultured with STING agonist-stimulated DCs. These findings collectively demonstrate that STING agonists are capable of both activating and inducing proliferation in CD8+ naïve T cells, supporting their potential role in immunotherapeutic applications.

**Figure 3.**
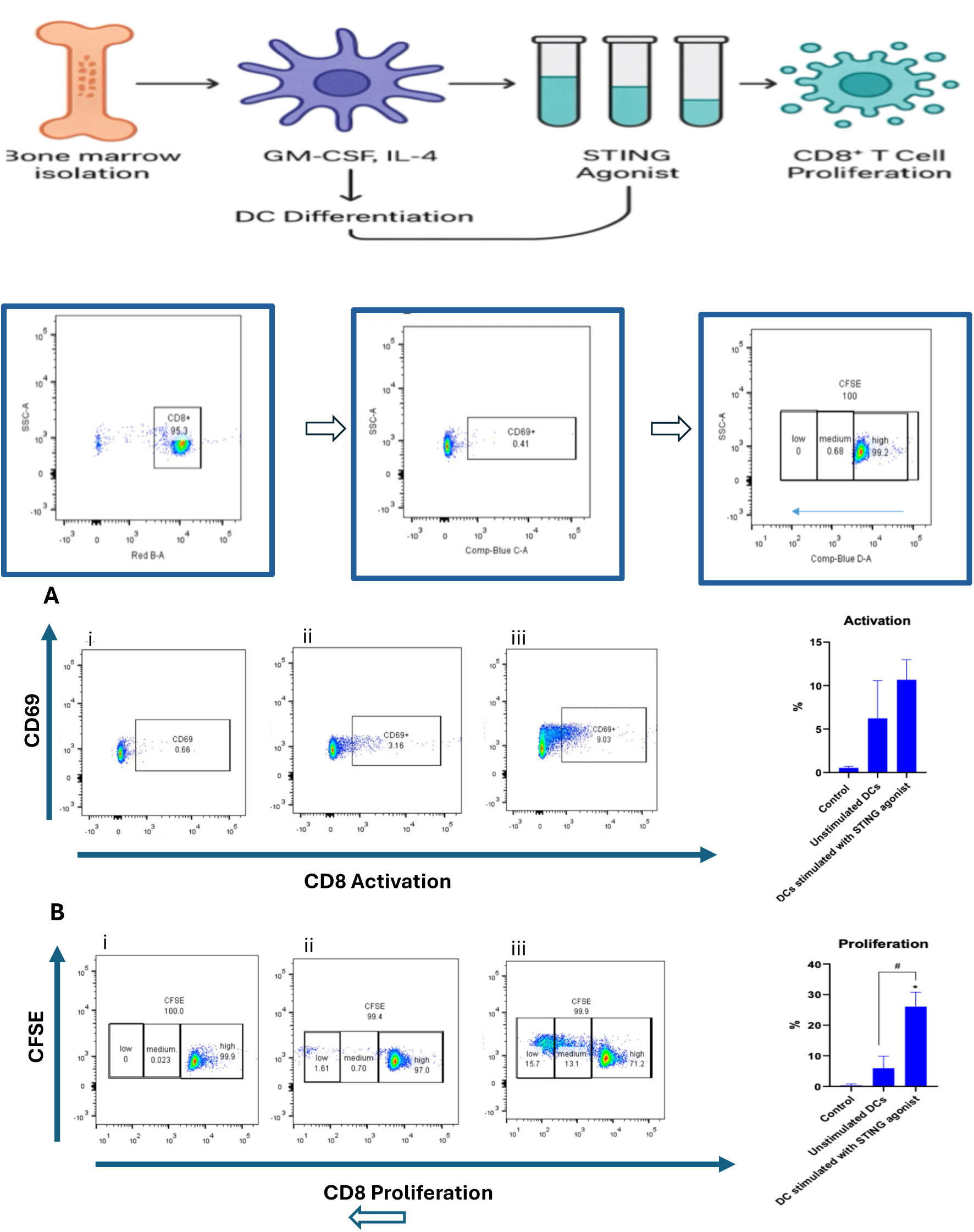
Activation and proliferation of CD8⁺ naïve T cells following co-culture with dendritic cells (DCs). Representative flow cytometry plots showing CD8⁺ T cell activation under the following conditions: (A) (i) CD8⁺ naïve T cells without any stimulation; (ii) CD8⁺ T cells co-cultured with unstimulated DCs; (iii) CD8⁺ T cells co-cultured with DCs stimulated with 5 µg/mL STING agonist. (B) CD8⁺ T cell proliferation assessed under the same conditions: (i) CD8⁺ naïve T cells without stimulation; (ii) T cells co-cultured with unstimulated DCs; (iii) T cells co-cultured with DCs stimulated with 5 µg/mL STING agonist. Data are presented as mean ± SD of two independent experimental replicates, and were analyzed using two-way ANOVA, with significance indicated by *p < 0.05 and #p < 0.05.

## 4. Discussion

In this study, we evaluated the ability of dendritic cells (DCs) stimulated with a STING agonist to activate and induce the proliferation of naïve CD8⁺ T cells. Our results demonstrate that STING pathway activation in DCs significantly enhances both the early activation and clonal expansion of CD8⁺ T cells. Specifically, we observed a marked upregulation of CD69, an early activation marker, on CD8⁺ T cells co-cultured with STING-stimulated DCs compared to those cultured with unstimulated controls. In parallel, CFSE-based proliferation assays revealed a substantial increase in cell division, indicating that STING activation not only initiates T cell activation but also sustains proliferative signaling necessary for effective immune responses.

These findings align with previous reports showing that STING activation in antigen-presenting cells can enhance T cell priming through upregulation of costimulatory molecules (e.g., CD80, CD86) and secretion of type I interferons and pro-inflammatory cytokines. By bolstering the immunostimulatory capacity of DCs, STING agonists may help overcome the immunosuppressive barriers commonly encountered in the tumor microenvironment and improve the efficacy of T cell–mediated immune responses.

Interestingly, we noted a moderate increase in CD69 expression in CD8⁺ T cells co-cultured with unstimulated DCs, suggesting that even in the absence of strong exogenous stimulation, DCs can provide a baseline level of T cell activation. This effect is likely mediated by tonic MHC-TCR interactions and/or low-level cytokine production. However, this basal activation was insufficient to drive significant T cell proliferation, emphasizing the necessity of robust costimulatory and cytokine signals for full T cell activation and expansion—signals that are clearly amplified upon STING engagement.

Our observation that STING-activated DCs promote not only activation but sustained proliferation of CD8⁺ T cells supports the idea that STING signaling reinforces both the initiation and maintenance phases of T cell responses. This has important implications for cancer immunotherapy, where T cell exhaustion, lack of effective antigen presentation, and immune evasion are major obstacles to success. By improving the immunogenic profile of DCs and enhancing their capacity to prime cytotoxic T lymphocytes (CTLs), STING agonists hold significant promise as adjuvants in cancer vaccines or combination immunotherapies.

Moreover, the ability to drive the expansion of antigen-specific T cells in vitro under defined conditions provides a valuable platform for preclinical evaluation of personalized immunotherapies. This strategy could be particularly useful for generating patient-specific T cell responses against tumor neoantigens or viral epitopes, thereby improving the precision and efficacy of treatment.

In conclusion, our findings underscore the potent immunostimulatory effects of STING pathway activation in dendritic cells, leading to enhanced activation and proliferation of CD8⁺ naïve T cells. These results support the continued development of STING agonists as adjuvant agents in immunotherapy and highlight the potential of integrating STING signaling into DC-based vaccine platforms. Future in vivo studies are warranted to evaluate the therapeutic efficacy, safety, and potential synergistic effects of STING agonists in cancer and infectious disease models.

## 5. Remarks

This study aimed to explore the role of GM-CSF and IL-4 in dendritic cell (DC) maturation and to investigate the contribution of DNA sensing mechanisms particularly the STING pathway on CD8+ T cell proliferation. Our findings highlight the importance of STING signaling in initiating effective T cell responses, specifically demonstrating that CD8+ T cells ^17^, which play a central role in tumor clearance, can be activated and proliferated via STING pathway engagement.

Given the inherent difficulty of maintaining and expanding T cells in vitro, future research should prioritize the optimization of T cell culture media to improve proliferation efficiency and cellular viability. Additionally, fine-tuning the co-culture conditions such as DC-to-T cell ratios and timing may enhance T cell activation and expansion outcomes.

To better understand the full landscape of CD8+ T cell activation, future studies should also investigate the involvement of various cytokines and chemokines that are secreted during DC–T cell interaction. Elucidating the individual contributions of activation signals—Signal 1 (antigen recognition via MHC-TCR) and Signal 2 (co-stimulatory molecule interactions)—may uncover potential bottlenecks or enhancement points in T cell priming.

Finally, these findings suggest promising implications for cancer immunotherapy. Exploring DNA-based antigens as potential cancer vaccine candidates could open new avenues for tumor-specific immune activation. Further in vivo validation and translational studies are necessary to assess the therapeutic efficacy and safety of such approaches in clinical settings.

## Conflicts of Interest

The authors declare that there is no conflict of interest regarding the publication of this article.

## Author Contributions

Busra Buyuk: Conceptualization, Methodology, Data Curation, Writing – Original Draft. Kaiming Ye: Supervision, Review & Editing.

## Acknowledgement

I thank Subhadra Jayaraman for her support with the experimental setup and assistance with the flow cytometry analysis.

## Notes

### Competing Interest Statement

The authors have declared that no competing interests exist.

### Summary of Updates

The figures in the previous PDF version were not clearly visible; therefore, I have re-uploaded the figures in a separate Word document to ensure better resolution and clarity. In addition, I have made minor corrections to a few inaccurate sentences in the Materials and Methods section.

